# A multispecies framework for modeling adaptive immunity and immunotherapy in cancer

**DOI:** 10.1101/2022.02.10.479985

**Authors:** Timothy Qi, Benjamin Vincent, Yanguang Cao

## Abstract

Predator-prey theory is commonly used to approximately describe tumor growth in the presence of selective pressure from the adaptive immune system. These interactions are mediated by the tumor immunopeptidome (what the tumor “shows” the body) and the T-cell receptor (TCR) repertoire (how well the body “sees” cancer cells). Importantly, both tumor and T cell populations are dynamic and in competition with each other and their fraternal lineages. In particular, the immunopeptidome comprises neoantigens which can be gained and lost throughout tumorigenesis and treatment. Heterogeneity in the immunopeptidome is predictive of poorer survival and response to T-cell dependent immunotherapy in some tumor types, suggesting the TCR repertoire is unable to support a fully polyclonal response against every neoantigen. Whether between-lineage competition among T cells plays a role, and in what contexts, is unknown; moreover, longitudinal TCR profiling studies are expensive and logistically complex to conduct. In silico models may offer an inexpensive way to interrogate these phenomena ex ante and deepen our understanding of the tumor-immune axis. Here, we construct and calibrate a predator-prey-like model to preclinical and clinical data to describe tumor growth and immunopeptidome diversification. Simultaneously, we model the expansion of neoantigen-specific T cell lineages and their consumption of both lineage-specific and shared, lineage-agnostic resources. This predator-prey-like framework accurately described clinically observed immunopeptidomes as well as correlates and consequences of response to immunotherapy, including immunoediting. The model was also suitable for exploring treatment of tumors with varying growth and mutation rates.

## INTRODUCTION

Cancer is a disease with parallels to predator-prey theory [1]. Control of prey (cancer cells) by predators (cytotoxic T cells, CTLs) depends upon the latter recognizing and killing the former. This process is mediated by the interaction between neoantigens on the surface of cancer cells and T cell receptors (TCRs) on the surface of CTLs. Neoantigens and TCRs that bind each other are called cognate; cognate interactions trigger CTLs to clonally expand, producing daughter cells that express the same TCR. At the molecular level, neoantigens arise from tumor-specific alterations to DNA or RNA and are bound by major histocompatibility complex (MHC) molecules for extracellular presentation to TCRs (Figure 1A) [2], [3]. The myriad neoantigens presented by a population of cancer cells, exclusive of self antigens, is referred to here as the immunopeptidome, while the universe of TCRs expressed by all CTLs in the body is referred to here as the TCR repertoire. Importantly, while one cancer cell can present a theoretically unlimited number of neoantigens, individual T cells express TCRs of only a single specificity.

**Figure 1.**
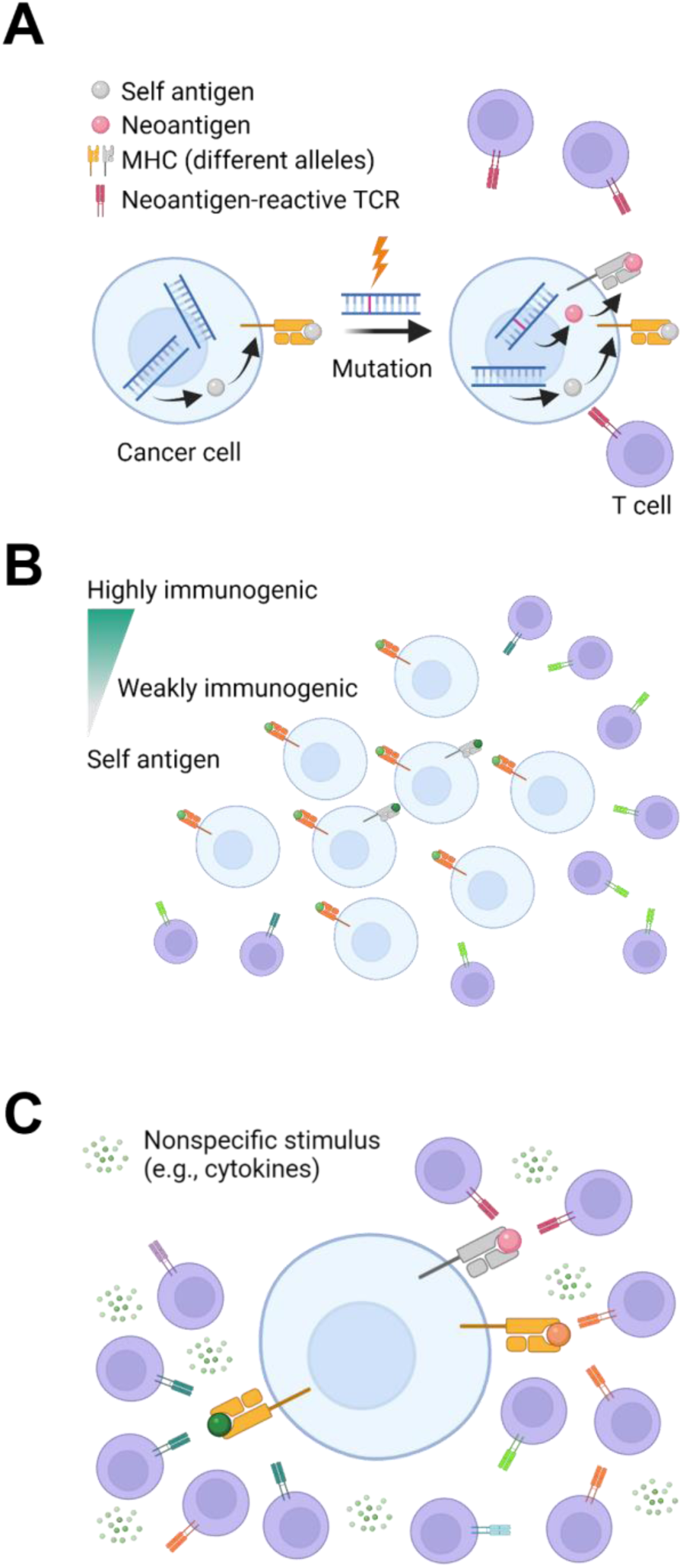
Intratumoral predator-prey dynamics depend on the immunopeptidome A. Schematic of neoantigen presentation. Neoantigens arriving from tumor-specific aberrations in DNA or RNA are presented on MHC molecules. CTLs with TCRs that recognize the neoantigen subsequently proliferate and attack the neoantigen-presenting cell. B. Schematic of immunodominance. Highly immunogenic neoantigens that are also subclonal can be shielded from CTLs due to the abundance of more clonal neoantigens. C. Schematic of interclonal competition. CTLs reactive against different neoantigens compete for the same pool of shared nonspecific stimulus, including cytokines and co-stimulation from antigen-presenting cells.

CTLs kill cancer cells that present their cognate neoantigen. The propensity of a neoantigen to cause this is referred to as its immunogenicity. Numerous factors influence immunogenicity, including several key parameters reported by the global community-based Tumor Neoantigen Selection Alliance [4]. These parameters comprise (1) MHC binding affinity, MHC binding stability, or duration of peptide-MHC interaction, (3) agretopicity, or the ratio of mutant to wild-type binding affinity, (4) RNA transcript abundance within the tumor, and (5) foreignness, or homology to epitopes of known pathogens. These parameters and their associated cutoffs associated with immunogenicity are referred to here as the TESLA criteria.

The heritability of both neoantigens and their cognate TCRs over cellular generations is important to consider when evaluating anti-neoantigen immunity. For one, clonal expansion of distinct cancer and T cell lineages is common. Tumors enriched in clonally expressed neoantigens are more highly infiltrated by CTLs, particularly those with cognate TCR clonotypes. Highly clonal neoantigen burden correlates with longer survival in patients treated with T cell-dependent cancer immunotherapies, such as immune checkpoint blockade (ICB) [5], [6]. Furthermore, a recent pan-cancer analysis found clonal tumor mutation burden (TMB), a measure closely related to clonal neoantigen burden, to be among the strongest predictors of response to ICB [7].

Neoantigen clonality plays a particularly important role in differentiating between intrinsic and apparent immunogenicity. When an antigen is subclonally expressed below a certain fraction, lineages presenting these antigens may not be efficiently eliminated by the immune system even if the antigen has high intrinsic immunogenicity (e.g., chicken ovalbumin antigen, a highly foreign model antigen commonly used in murine studies) (Figure 1B) [8], [9]. Mechanisms for subclonal neoantigen include not only de novo generation, but also through (epi)genomic silencing of extant neoantigens and/or their associated MHC alleles [10], [11]. In humans, studies across multiple cancer types have found that among hundreds of putative neoantigens identified by bioinformatics pipelines, very few (generally < 4) are bona fide immunogenic when tested in ex vivo assays; moreover, those that are immunogenic tend also to be clonal [4], [5], [12]–[15].

It is unclear why the TCR repertoire does not support a fully polyclonal response against every neoantigen. We speculate this limitation could be due to the spatiotemporally restricted availability of resources and nutrients critical to CTL survival and proliferation [16]. For example, cytokines, co-stimulatory signals from antigen-presenting cells, and other factors beyond direct antigen-mediated signaling through TCRs are a shared resource among CTL lineages (Figure 1C) [16], [17]. The asymmetric and possibly contemporaneous presentation of multiple neoantigens might exhaust these nonspecific stimuli, leading to an inadequate immune response to subclonal neoantigens. In humans, the clonal expansion of antigen-reactive CTL lineages can indeed suppress the expansion of other CTL lineages, a phenomenon called immunodominance that is well characterized within virology [18]–[20]. In cancer, ICB has been widely reported to affect the diversity of the CTL TCR repertoire, although whether it biases CTL expansion toward polyclonal or oligoclonal states appears to vary by ICB target and tumor histology [21]. However, given the shared nature of nonspecific stimuli, highly clonal tumors might be more susceptible to CTL-based immunotherapies due to reduced interclonal competition for nonspecific stimuli as compared to highly subclonal tumors rife with lower-quality, “distracting” neoantigens [16].

Neoantigen quantity is therefore critical to consider alongside immunogenicity for immune recognition and ICB treatment efficacy [15], [22]. However, the quantitative relationship between immunopeptidome diversity, CTL diversity, and response to ICB is open to further characterization. In this work, we construct a mathematical model of nonlinear predator-prey-like dynamics to describe the relationship between immunopeptidome diversity and CTL responses while accounting for interclonal CTL competition for nonspecific stimuli. Tumor growth and immunopeptidomics are modeled and calibrated with experimental data from two different species and three types of cancer, including an orthotopic murine model of bladder cancer, a murine model of colorectal cancer, a 100-patient cohort of NSCLC patients who underwent extensive immunopeptidome profiling [10], [11], [23]–[25]. Simulated NSCLC-like immunopeptidomes and TCR repertoires are cross-validated on an independent cohort [26] and subject to a local sensitivity analysis. Finally, tumor response dynamics in the NSCLC-like cohort, as well as in two additional cohorts resembling other tumor types, are simulated under ICB treatment to evaluate pre-treatment biomarkers of response within the immunopeptidome and the TCR repertoire.

## METHODS

### Tumor growth dynamics

Tumors grew according to a modified logistic growth model, where *N* is the number of cells in the tumor, *k_g_* is the crude birth rate, K is the population carrying capacity, *k_d,i_* is the lineage-specific death rate (see *Lineage-specific killing*), and *j* is number of unique cancer cell lineages in the tumor:

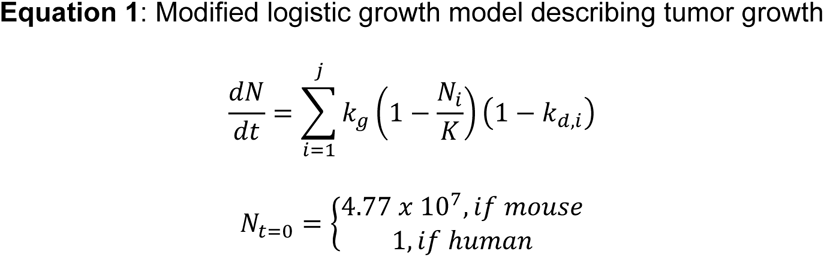

All cancer cell lineages were assigned the same birth rate and shared population-level carrying capacity. Death rates varied by lineage and depended on the number of CTLs cognate to the lineage’s neoantigens (see respective sections on *CTL clonal dynamics* and *Neoantigen features*). Cancer cell numbers were converted into tumor diameters assuming spherical tumors and an approximate cell volume of 4 pL [27].

During mouse simulations, tumors were initiated from a single lineage ∼4.77 x 10^7^ cells, corresponding to an approximate volume of 200 mm^3^ at which tumor measurements began in [23], [28]. During human simulations, tumors were initiated from a single founder cell with a gamma-distributed number of neoantigens [10]. Furthermore, the founder cell was assigned a 14% chance of having an MHC loss event [11] in one of its three MHC alleles. Further details on neoantigen immunogenicity, expression, and MHC presentation are provided in the *Neoantigen features* section. Parameter values, definitions, and their sources are provided in Table S1.

### CTL clonal dynamics

At baseline, we assumed a tumor-ignorant and nonspecific CTL population equal to the steady-state population of T cells *CTL_ss_* in either mice or humans (Table S1). With the number of unique neoantigens *k* known from either a founder lineage (mice) or cell (humans), we then created *k* tumor-reactive CTL lineages (Figure 2). For simplicity, we assume TCRs are specific to a single neoantigen, which aligns with the high specificity of TCRs despite their considerable cross-reactive potential [29]. For each neoantigen among *k* neoantigens, a CTL lineage of 2^n^ cells was created, where n is a Poisson-distributed number *n* ∼ *P(µ)*. This represents the random number of divisions the founder CTL cell underwent prior to thymic efflux [30]. The remaining nonspecific CTLs were lumped into a tumor-ignorant, nonspecific population with a starting cell number equal to *CTL_ss_* minus the total number of neoantigen-reactive CTLs.

**Figure 2.**
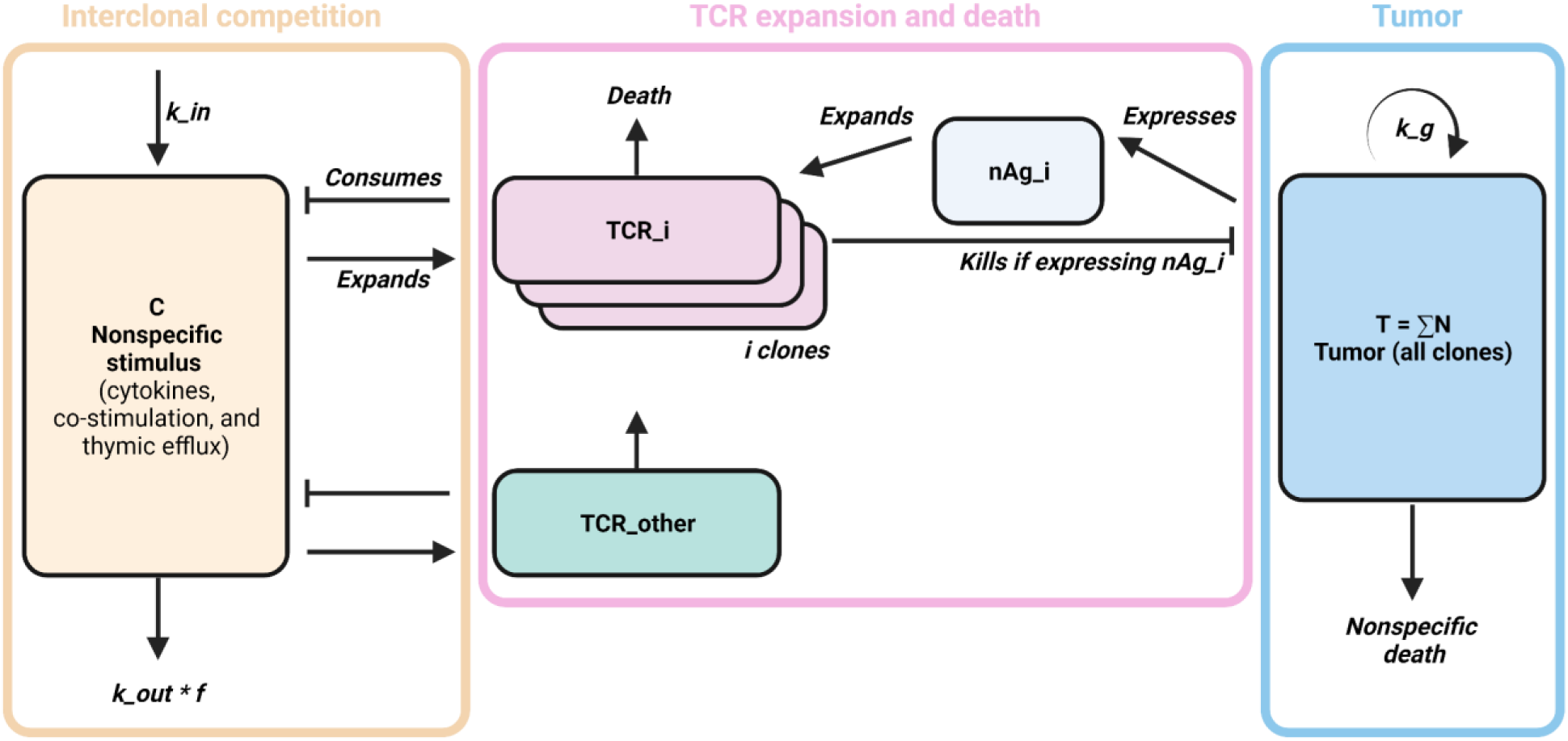
Distinct tumor-reactive CTL lineages were modeled to capture immunopeptidome interactions A. Schematic of the TCR-specific CTL lineage model. Cancer cells present neoantigens, providing pro-proliferative stimulus to neoantigen-specific populations of CTLs. A fraction of cancer cells dies to CTL-mediated killing, while the remaining fraction dies to unrelated causes. In parallel, CTL expansion depends on sufficient nonspecific stimulus, which is consumed by all tumor-reactive CTL lineages as well as a pooled lineage representing all tumor-ignorant CTLs. A fraction of nonspecific stimulus turns over irrespective of CTL consumption.

Tumor-specific CTL lineages grew or shrank according to the quantity of their cognate antigenic stimulus and the availability of shared nonspecific stimuli, where *CTL_k_* is the *k*th CTL lineage reactive to the *k*th neoantigen, *T_d_* is its baseline turnover rate, *C* is the abundance of nonspecific stimulus, *S_k_* is the immunogenicity-weighted number of cancer cells expressing the *k*th neoantigen (see *Neoantigen features*), and *T_a_* is a scaling factor relating the rate of *CTL_k_* expansion per cancer cell expressing the *k*th neoantigen:

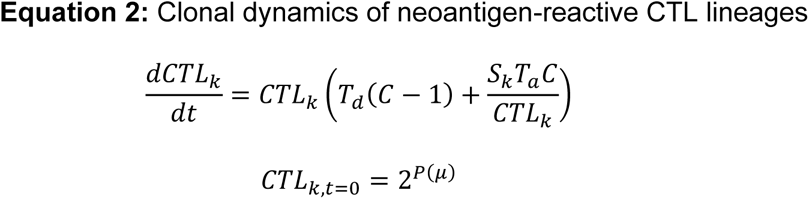

Nonspecific CTLs presented with no antigen stimulus grew or shrank according only to the availability of shared nonspecific stimuli, where *NSCTL* is the number of non-specific CTLs:

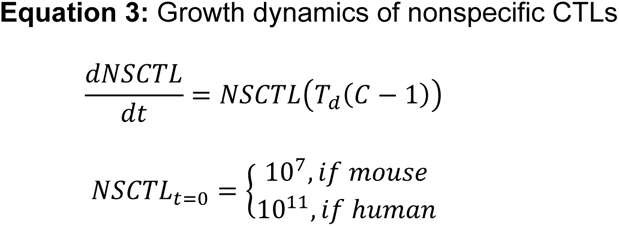

Nonspecific stimulus *C* was consumed by T cells and non-T cells and replenished at rate *k_in_*. The fraction of *C* consumed by non-T cells was *f_out_*. *C* therefore decreased when the total number of nonspecific and tumor-reactive CTLs was greater than *CTL_ss_* and increased when it was less than *CTL_ss_*:

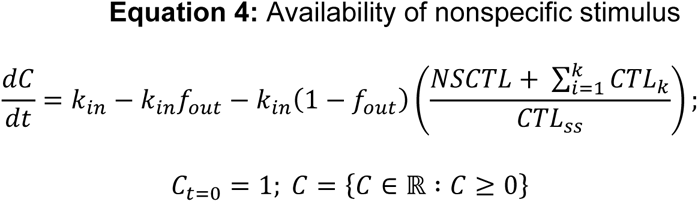

Shannon entropy among the tumor-reactive CTL lineages was calculated using MATLAB’s *wentropy* function.

### Neoantigen features

Each neoantigen was assigned a non-negative real number *R* between 0 and 1 to describe its immunogenicity. In the mouse simulations, *R* for each of the 34 neoantigens identified in BBN963 cells in [23] was assigned by scaling the baseline-corrected ELISpot scores to the range [0, 1].

In the human simulations, *R* was assigned for each neoantigen in the founder cell by first sampling values for each of the TESLA criteria. Neoantigens were first assigned random values for MHC binding affinity, MHC binding stability, agretopicity, transcript abundance, and foreignness drawn from distributions defined by 146 NSCLC-derived peptides (Figure S1) [4]. To achieve this, we fit an exponential distribution to transcript abundance and a zero- and one-inflated binomial distribution to foreignness. To account for correlation among MHC binding affinity, MHC binding stability, and agretopicity values, we fit a Gaussian copula to log-transformed values using MATLAB’s *copulafit* function. Parameters associated with these fits are provided in Table S1. Transcript abundance was discarded due to conflation with clonality and the remaining four parameters were assessed against the following TESLA criteria: MHC binding affinity < 34 nM, MHC binding stability > 1.4 hours, agretopicity > 0.1, and foreignness < 10^-16^. Neoantigens that failed to satisfy these cutoffs were assigned *R* = 0, or no immunogenicity. Neoantigens that passed these cutoffs were assigned a value for *R* from an exponential distribution:

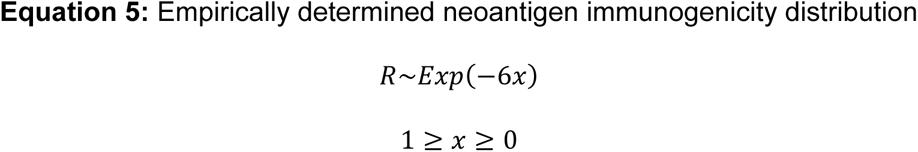

In addition to intrinsic immunogenicity, each neoantigen was also assigned values to indicate its expression status and MHC binding partner. In all cases, neoantigens were assumed to be expressed at baseline. Murine simulations assumed each neoantigen could bind to one of two MHC alleles (H2-Db and H2-Kb), both of which were intact and expressed at baseline. Human simulations assumed neoantigen binding to one of three MHC alleles (HLA-A, HLA-B, and HLA-C), all of which were expressed at baseline, except in ∼14% of tumors where loss of one of three alleles at random conferred clonal MHC loss [11]. Thus, inclusion of a neoantigen in its parent cell’s immunopeptidome depended upon its own expression status as well as the intactness of its binding MHC allele (Figure S1).

*S_k_*, the immunogenicity-weighted number of cancer cells with neoantigen *k* in their immunopeptidome, was used to stimulate the proliferation of *CTL_k_* per *CTL clonal dynamics*. It was calculated by multiplying the intrinsic immunogenicity of a neoantigen *k* by the number of cells whose immunopeptidomes include it, where *E_k_* is the lineage’s expression status of neoantigen *k*, *MHC_k_* is the lineage’s expression status of the MHC binding partner of neoantigen *k*, and *R_k_* is the intrinsic immunogenicity of neoantigen *k*:

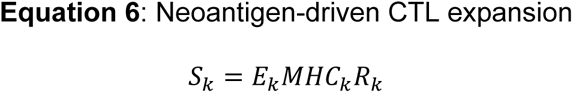

In mouse simulations, cancer cell lineage’s immunopeptidomes were held constant within the short simulated timeframe due to the lack of data to inform event rates. In human simulations, stochastic modifications to a cancer cell lineage’s immunopeptidome were allowed to occur. These events were one of three types: (1) neoantigen gain, (2) neoantigen loss, and MHC allele loss. Neoantigen gain events, representing the generation of new and heritable neoantigens within the DNA or RNA, involved drawing TESLA parameters and assigning *R* immunogenicity values for each new neoantigen as described above. Neoantigen loss events, representing the genetic, epigenetic, transcriptomic, or metabolic suppression of a neoantigen, were represented as an irreversible change in the neoantigen’s expression from 1 to 0; functionally, this removed the neoantigen from the lineage’s immunopeptidome and reduced its *S_k_* to 0. MHC loss events, analogous to neoantigen loss events for MHC alleles, were represented as an irreversible change in one of the three MHC alleles’ expression from 1 to 0. Functionally, this removed all neoantigens presented by that MHC allele from the lineage’s immunopeptidome and reduced their *S_k_* to 0. Lineages that experienced one of these stochastic events were deducted 1 cell from their population size. An identical lineage save the stochastic event was then created at N=1.

### Lineage-specific killing

Each unique clone in the tumor was killed at an independent and saturable rate calculated from the contents of its immunopeptidome and the abundance of cognate, reactive CTLs. Here, *k_d,i_* is a dimensionless value used to modify the logistic growth described in Equation 1; *f_die_* is the fraction of cell death not attributed to CTL killing, *k_max_* and *k_kill_* are shape parameters describing the saturable killing function, and *j* is the number of unique neoantigens harbored by the lineage:

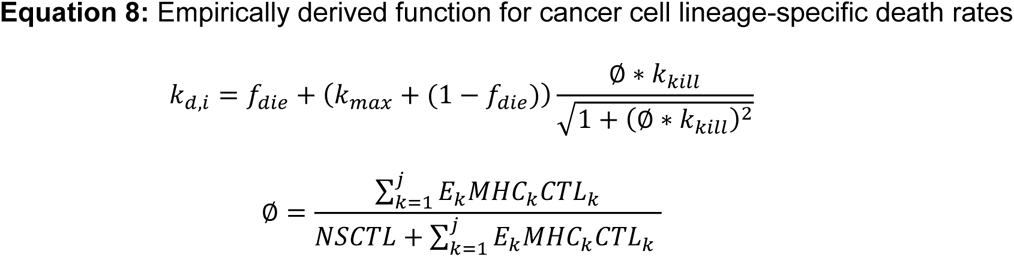

Thus, the total CTLs reactive against *j* neoantigens is summed to calculate antigen-specific killing. *CTL_k_* is only counted if the *k*th neoantigen and its MHC are expressed (*E_k_* and *MHC_k_* = 1).

### Simulated tumor growth in mice

BBN963 orthotopic bladder cancer cells were previously established by chronic exposure of C57BL/6 mice to 0.05% N-Butyl-N-(4-hydroxybutyl) nitrosamine [23]. Neoantigens were predicted computationally and screened for bona fide immunogenicity in an ex vivo ELISpot assay (Figure S2A). Additional details can be found in [20]. BBN963 cells were implanted in immunocompetent C57BL/6 or immunodeficient NSG mice [26]. Tumor volumes were measured weekly for five weeks once the tumors reached 200 mm^3^. Tumor growth volumes were digitized using WebPlotDigitizer [31] and used to calibrate mouse-specific model parameters in Table S1.

An independent dataset describing the growth dynamics of MC38 murine colon cancer cells in immunodeficient and immunocompetent mice were used to validate the mouse model. MC38 tumors were subcutaneously established in immunocompetent C57BL/6 or immunodeficient RAG1 KO mice [25]. Whole exome and RNA sequencing were used to identify 489 expressed neoantigens. In the absence of reported MHC allele specificity and ex vivo-validated immunogenicity, each neoantigen was assigned a random for MHC allele specificity and immunogenicity from the distribution reported for BBN963 cells [23]. Tumor growth in both immunodeficient and immunocompetent mice was successfully recapitulated by using the same parameters as BBN963 cells, except growth rate, which was increased 50% (Figure S2B-D).

### Simulated tumor growth in humans

The TRACERx 100 cohort [24] contained 100 patients with varying stages of NSCLC: 26 stage I, 36 stage IB, 13 stage IIA, and 25 stage IIB/IIIA/IIIB. These patients underwent multiregion biopsy for extensive immunopeptidome profiling [10], [11]. Tumor diameters were estimated for each stage using conventional NSCLC staging criteria [32]. Clonal and subclonal neoantigen burden digitized from [10] with WebPlotDigitizer, as well as rates of neoantigen expression loss and MHC clonal/subclonal loss of heterozygosity reported in these studies, were used to calibrate simulated immunopeptidomes. In total, 100 tumors were simulated to target diameters mirroring the TRACERx 100 cohort. Clonal and subclonal neoantigen burden for each tumor was then calculated by determining the number of neoantigens present in 100% or < 100% of lineage-specific immunopeptidomes.

### Human immunopeptidome cross-validation and local sensitivity analysis

Simulated immunopeptidomes were evaluated against an independent cohort of 12 patients [26] (Figure S3). Putative neoantigens from single-region biopsies collected from 12 patients prior to ICB treatment were assessed. Total neoantigens were calculated by filtering for positive netCTL classification. To apply the TESLA criteria, we calculated agretopicity by taking the ratio of somatic and wild-type MHC binding affinity. For predictions containing somatic peptides of identical sequences but different predicted MHC binding alleles, the more strongly binding prediction was used. Binding stabilities were calculated with NetMHCStabPan using default parameters [33]. The number of neoantigens that passed three of five the TESLA criteria (MHC binding affinity < 34 nM, MHC binding stability > 1.4 hours, and agretopicity > 0.1) was multiplied, in the absence of reported foreignness, by a correction factor equal to the ratio of neoantigens in the TRACERx 100 [10] cohort passing the MHC binding affinity, stability, and agretopicity criteria that also did or did not pass the foreignness cutoff of < 10^-16^. This reduced the number of predicted neoantigens in the immunopeptidome by 80.6%. Neoantigen clonality was not assessed due to potential sampling biases intrinsic to single-region biopsies. For each of the seven discrete tumor stages, tumor diameters were estimated for each stage using conventional NSCLC staging criteria [32]. A cohort of 40 tumors was simulated for each of the seven diameters. The simulated and experimentally reported immunopeptidomes are shown in Figure S3.

Local sensitivity analysis on the simulated TRACERx 100 cohort was performed by re-simulating each tumor after varying one estimated parameter up or down 20%. This was repeated for each estimated parameter in the model; the results are shown in Figure S4.

### Exploratory simulations of ICB treatment and other tumor histologies

A cohort of 100 tumors was simulated until a target population roughly equivalent to a ∼5 cm diameter tumor. ICB treatment of varying strength was implemented to increase the sensitivity of CTLs to antigenic stimuli as well as the population-level carrying capacity, as below (where *ICB* is a positive real number representing simulated treatment strength):

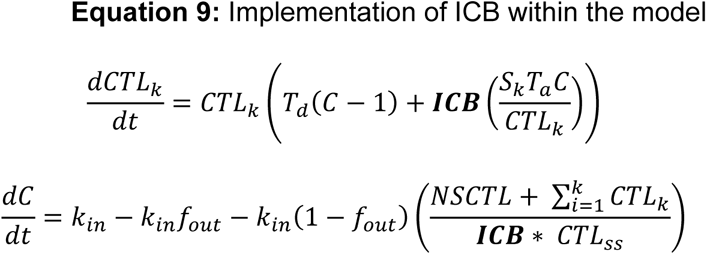

To emulate renal cell carcinoma (RCC)-like tumors, a cohort of 100 tumors with 50% lower growth rate and 90% lower neoantigen gain, neoantigen loss, and MHC loss rates was simulated. To emulate microsatellite instability-high colorectal cancer (MSI)-like tumors, a cohort of 100 tumors with 100% higher growth rate and 200% higher clonal neoantigen burden was simulated. ICB treatment was simulated as above.

### Statistical analyses

All simulations and statistical analyses were performed in MATLAB R2020b. Pearson’s and Spearman’s ρ were used as appropriate to assess linear correlation among continuous or ranked variables. Continuous variables in the local sensitivity were assessed for statistical significance at α = 0.05 using one-way ANOVA and a post-hoc pairwise Student’s t-test, with Bonferroni correction for multiplicity. In cases of post-hoc multiple hypothesis testing, a Bonferroni correction was applied. Details of specific statistical analyses are provided in their corresponding figure captions.

## RESULTS

### Model recapitulates tumor growth in immunocompetent and immunodeficient mice

BBN963 orthotopic bladder cancer cells generated in [23] were implanted in immunocompetent C57BL/6 and immunodeficient NSG mice [28] (Figure 3A). Thirty-four neoantigens putatively expressed by the cells were identified and tested for immunogenicity ex vivo [28] (see Methods). We used the results of these immunogenicity assays to scale in silico immunogenicity and assign one of two murine MHC alleles for presentation (Figure 3B). Five-week simulations of tumor growth in the absence of CTL expansion and CTL-mediated killing accurately reproduced BBN963 growth dynamics in NSG mice (Figure 3C). Similarly, five-week simulations with CTL expansion and cytotoxicity restored accurately reproduced BBN963 growth dynamics in C57BL/6 mice (Figure 3D).

**Figure 3.**
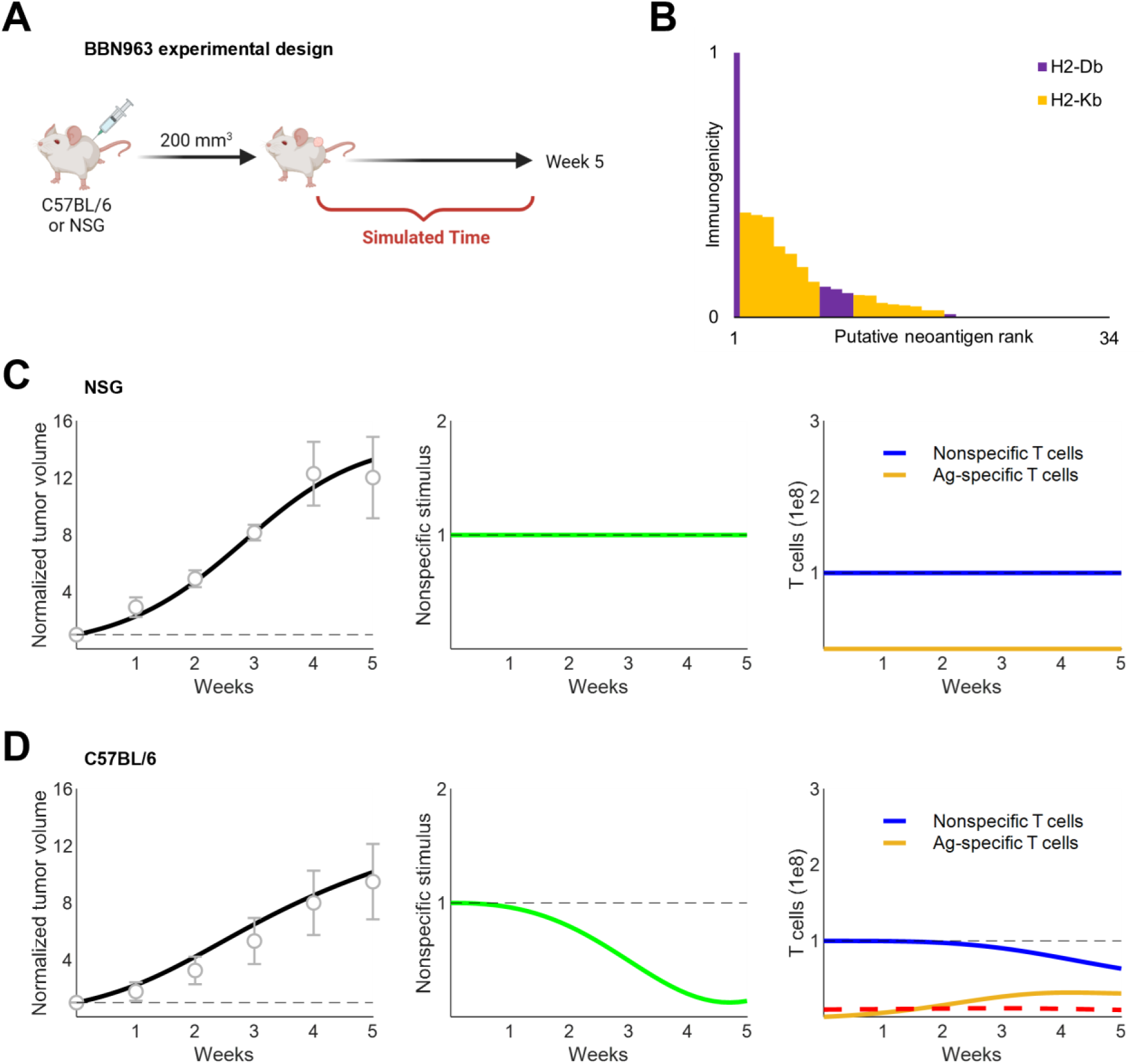
Murine tumor growth dynamics were recapitulated in divergent immune contexts A. NSG and C57BL/6 mice were inoculated with subcutaneous BBN963 tumors. Tumor volumes were measured over five weeks [28]. B. Saito et al. identified putative neoantigens expressed by BBN963 and the MHC alleles predicted to present them (H2-Db, H2-Kb). Baseline-corrected ELISpot scores were used to scale neoantigen immunogenicity to the range [0, 1]. C. Simulated tumor volumes (left, black) normalized to baseline volume (dashed gray line) compared against growth dynamics in NSG mice reported by Truong et al. (gray circles and error bars). Nonspecific stimulus (center) and CTL populations (right) are shown. Dashed lines, 1. D. Simulated tumor volumes (left, black) normalized to baseline volume (dashed gray line) compared against growth dynamics in C57BL/6 mice reported by Truong et al. (gray circles and error bars). Nonspecific stimulus (center) and CTL populations (right) are shown. Black dashed lines, 1; Red dashed lines, approximate total abundance of 3 Ag-specific T cell lineages observed in mice [28]

To test model fidelity against an independent dataset, we used the growth dynamics of MC38 murine colon cancer cells in immunodeficient and immunocompetent mice reported by Efremova et al. [25]. Assuming the same distribution of neoantigen immunogenicity as BBN963 cells and changing no parameters except growth rate, the model accurately captured MC38 tumor growth in both immunocompetent and immunodeficient mice (Figure S2B-D). These preliminary results suggested the generalizability of our predator-prey-like framework and motivated further simulation of human tumor-immune dynamics.

### Model recapitulates the tumor immunopeptidome of NSCLC patients

We substituted human parameter values for steady-state T cell abundance and lifespan from the mouse model, while also re-estimating tumor carrying capacity and T cell sensitivity to antigenic stimulus. No other parameters related to tumor or T cell dynamics were changed (Table S1). This enabled successful simulation of the growth and immunopeptidome of 100 human NSCLC tumors (Figure 4A) [24]. Tumor growth was simulated until target population sizes approximating the tumor diameters of TRACERx 100 NSCLC patients were reached. In contrast to the murine simulations, cancer cells in the human simulations were allowed to stochastically gain new neoantigens, lose expression of existing neoantigens, and lose MHC expression (see Methods, Figure S2A) [10], [11]. Simulations recapitulated the distribution of clonal, subclonal, and total neoantigen burden within this cohort (Figure 4B).

**Figure 4.**
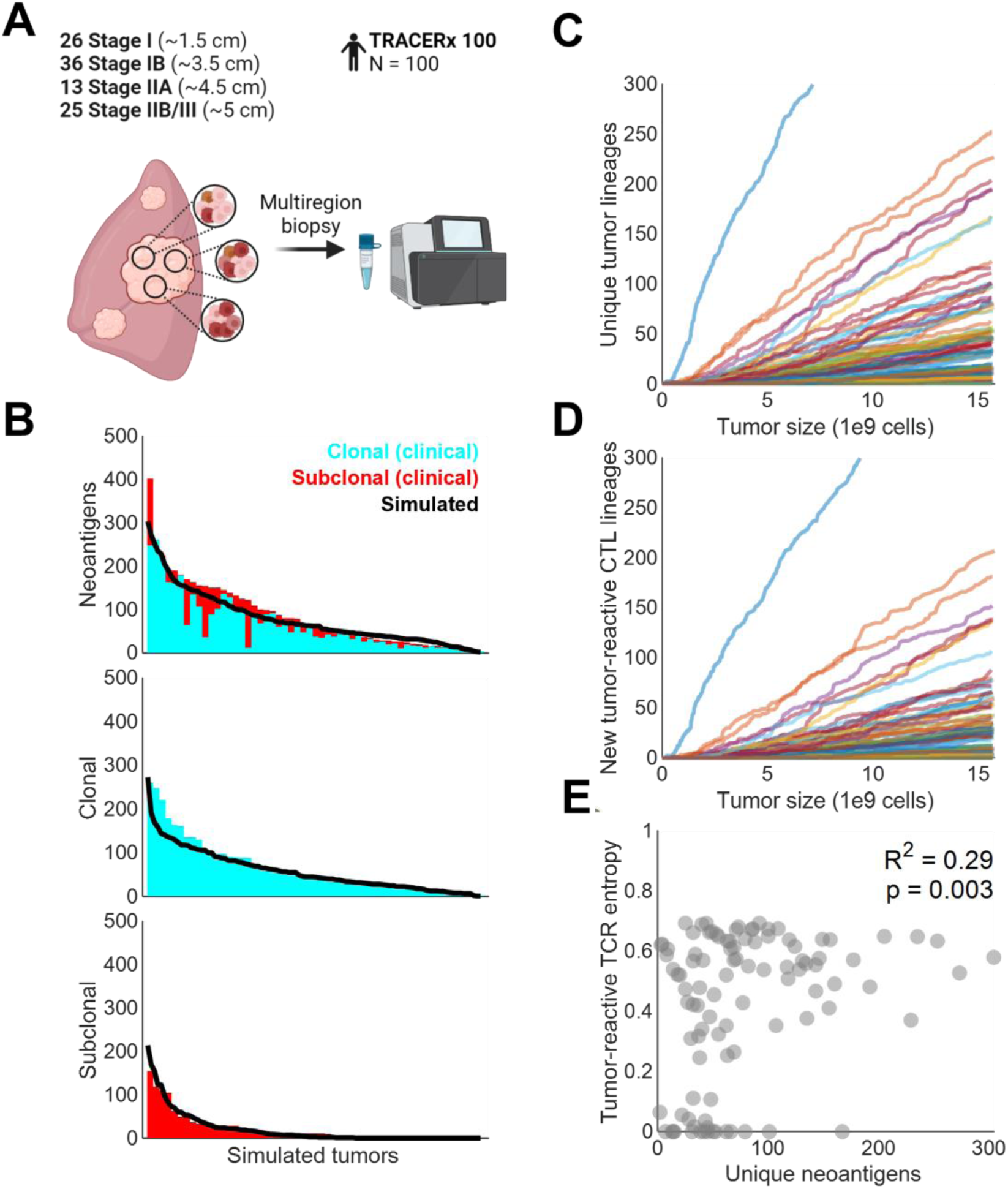
Human NSCLC immunopeptidomes were recapitulated from clinical data A. Schematic of multiregion biopsies collected in the TRACERx 100 cohort. 100 NSCLC tumors of various stages were sampled, corresponding to target diameters of 1.5 cm (stage IA), 3.5 cm (stage IB), 4.5 cm (stage IIA), and 5 cm (stage IIB/III). B. Simulated neoantigen burden compared against clonal and subclonal neoantigen burden digitized from [10]. C. Emergence of unique tumor lineages over the course of simulated tumor growth. D. Emergence of new tumor-reactive CTL lineages over the course of simulated tumor growth. E. Pearson’s ρ correlation between unique neoantigen number and tumor-reactive CTL Shannon entropy in the simulated TRACERx 100 cohort. Excludes one tumor with > 600 neoantigens for visualization purposes.

Interestingly, we observed substantial tumor and CTL diversification over time, as depicted in Figures 4C-D. The number of unique tumor lineages was consistently higher than the number of newly tumor-reactive CTL lineages due to lineage branching events caused by neoantigen and MHC loss. In these cases, a new cancer lineage was formed without introducing a new neoantigen and its cognate predator CTL. There was also a mild positive correlation between the number of unique neoantigens and CTL diversity (Figure 4E).

### Simulated ICB highlights pretreatment clonal neoantigen burden as a biomarkers of response

Prolonged antigen exposure results in CTL exhaustion, diminishing their proliferative and cytolytic capabilities. This process is accelerated by checkpoint molecules, many of which are overexpressed in the tumor microenvironment. ICB blocks these checkpoint molecules, reinvigorating the immune response and promoting the expansion of a select subset of CTL lineages (Figure 5A) [34]. To simulate ICB of late-stage NSCLC, we first constructed a 100- tumor virtual cohort comprising tumors of at least ∼5 cm in diameter. This roughly corresponds to the advanced stage of NSCLC for which ICB is indicated. Similar to the TRACERx 100 virtual cohort, we observed a mild positive correlation between neoantigen burden and CTL diversity prior to treatment (Figure 5B).

**Figure 5.**
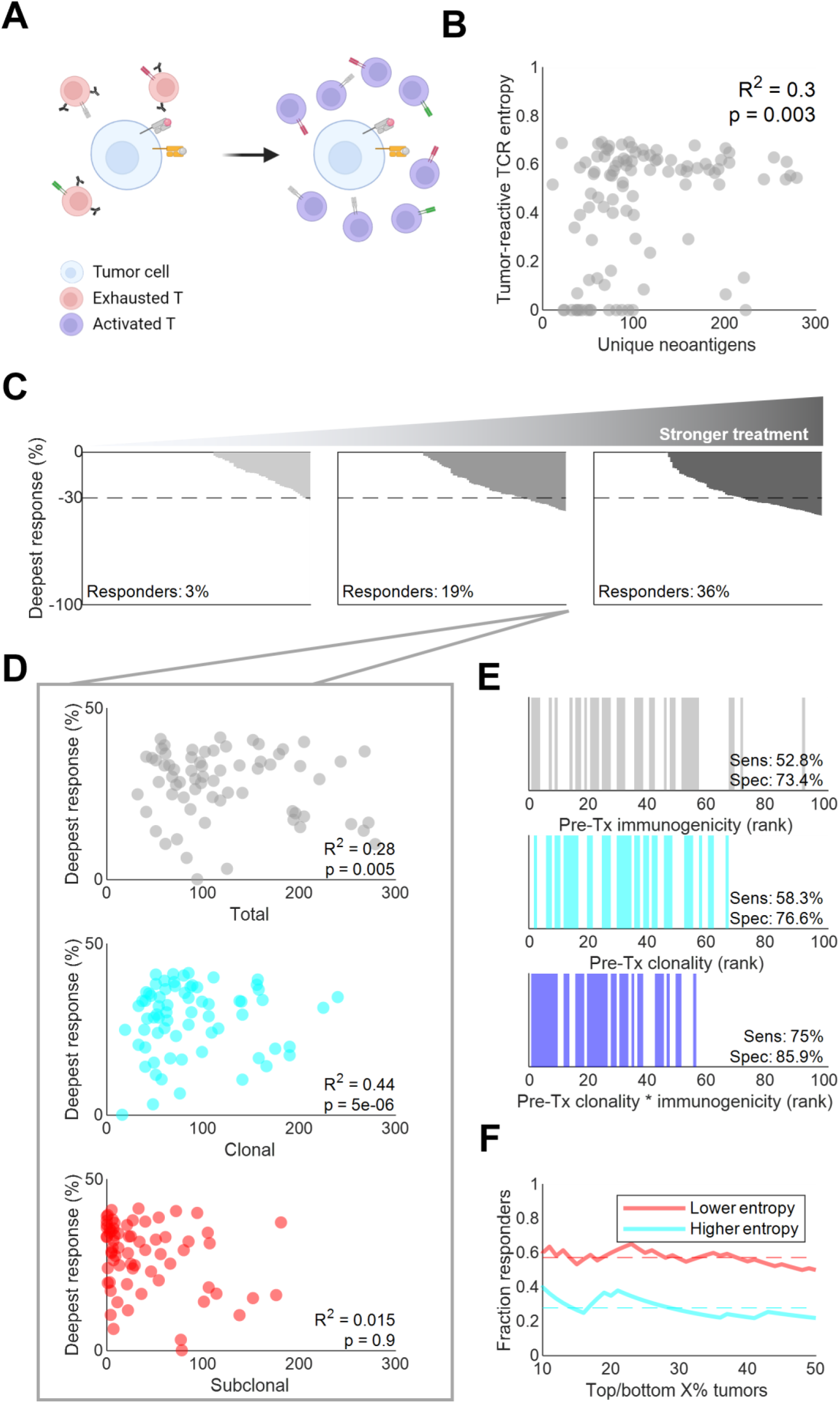
Pretreatment tumor-immune status correlated with response to immune checkpoint blockade A. Schematic of the proposed mechanism of action of PD-1 blockade, a common ICB. B. Pearson’s ρ correlation between unique neoantigen number and tumor-reactive CTL Shannon entropy in the simulated 100-tumor cohort prior to treatment. Excludes one tumor with > 600 neoantigens for visualization purposes. C. Maximum tumor shrinkage under increasing levels of treatment strength: 2- (left), 6- (center), and 10- (center) fold increases in sensitivity to antigen stimulus and systemic carrying capacity were simulated. Each bar corresponds to a tumor. The dashed line at corresponds to the 30% or greater tumor shrinkage required to classify as partial response per RECIST 1.1 criteria [35]. D. Pearson’s ρ correlation between response and pretreatment total, clonal, and subclonal neoantigen burden. Excludes one tumor with > 600 neoantigens for visualization purposes. E. Response status (colored bars) relative to rank 1 neoantigen immunogenicity (top), clonality (middle), and clonality-weighted immunogenicity (bottom). Sensitivity and specificity were calculated from the correctly classified fraction of responders (N=36) and non-responders (N=64), respectively. F. Fraction of responses among the X% most and least entropic immunopeptidomes, X being indicated by the X axis value. Entropy refers to Shannon entropy. Dashed lines, means.

Various strengths of ICB treatment were explored within the model by simulating tumor growth for one year in the presence of 2-, 6-, and 10-fold increases in CTL sensitivity to antigen stimulus with concurrent 2-, 6-, and 10-fold increases in steady-state CTL carrying capacity. As expected, the proportion of patients with maximal tumor shrinkage of at least 30% – “responders” – increased with treatment strength [35]. Notably, response rates in the 10-fold treatment simulation approached levels observed for ICB in Phase 3 clinical trials (Figure 5C) [36].

Pretreatment neoantigen burden correlated positively with deeper responses, in line with previous reports. (Figure 5D) [5]. Interestingly, this was driven primarily by clonal neoantigen burden. No significant relationship was found between response depth and the number of unique subclonal neoantigens. We also examined whether the immunogenicity or clonality of the most immunogenic neoantigen (i.e., rank 1) could predict response. In this case, clonality outperformed immunogenicity in predictive sensitivity and specificity; however, both were outperformed by the product of the two, termed clonality-weighted immunogenicity (Figure 5E). Responding tumors also had consistently lower Shannon entropy in their immunopeptidomes, corresponding to more homogeneous sets of neoantigens (Figure 5F). These data support the importance of neoantigen clonality and highlight intratumoral heterogeneity as a key detractor from ICB response [5], [37].

### CTL dynamics correlate with ICB response and immunoediting

While responders tended to have greater pretreatment CTL diversity, further CTL diversification induced by ICB treatment strongly correlated with shallower responses (Figure 6A-B). Instead, expansion and maintenance of CTL lineages that were highly abundant before ICB initiation tended to elicit more profound responses (Figure 6C) [21], [38], [39]. Responders also had more stable pools of nonspecific stimulus during treatment (Figure 6C). These findings support the notion that prognoses are most favorable when ICB induces focused, oligoclonal responses against highly clonal neoantigens, at least for certain tumor types [21], [40].

**Figure 6.**
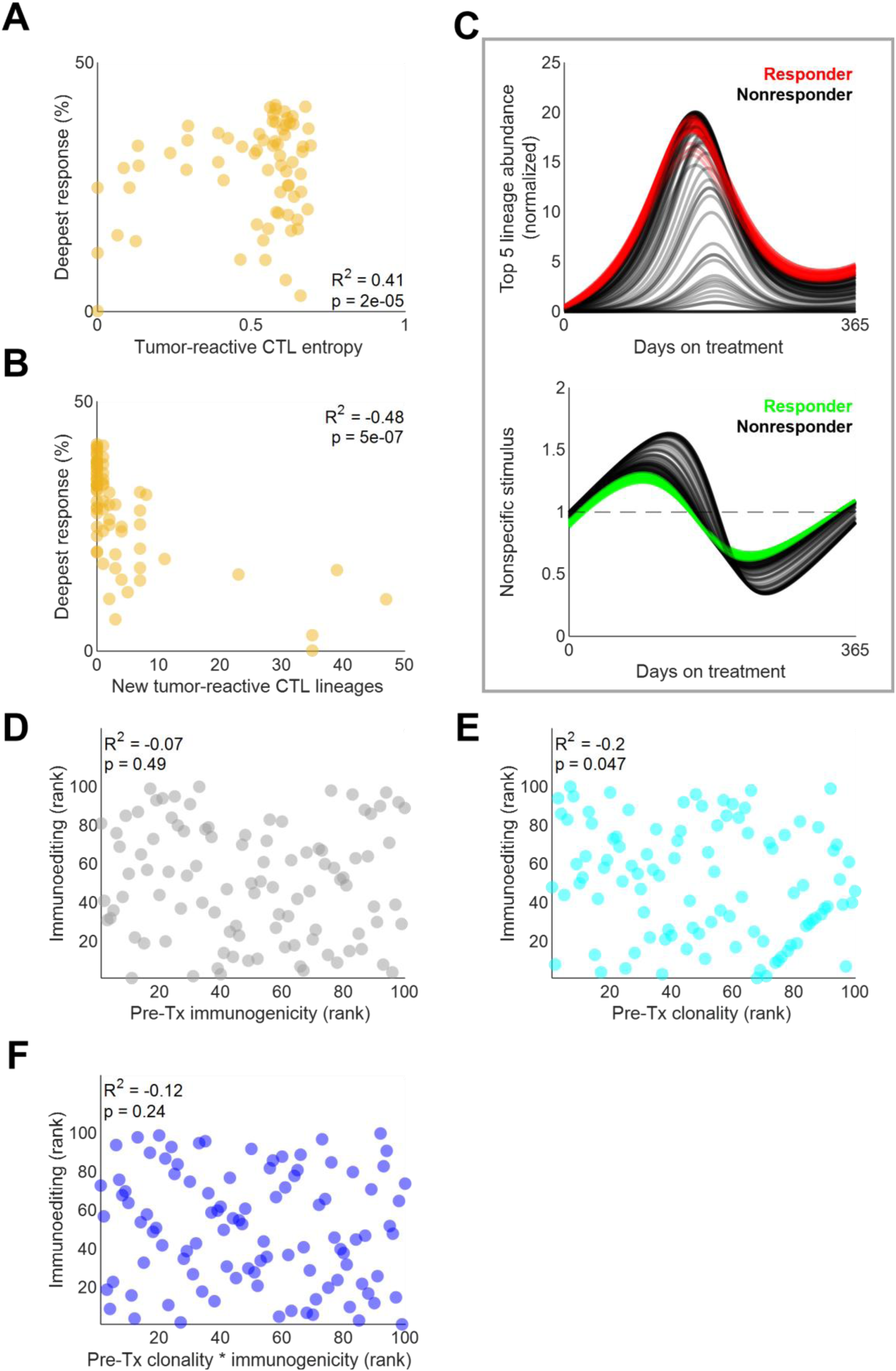
Immunopeptidome and CTL dynamics correlated with ICB response A. Pearson’s ρ correlation between response and pretreatment tumor-reactive CTL Shannon entropy. B. Pearson’s ρ correlation between response and new tumor-reactive CTL lineages that emerged upon simulated ICB. C. Summed abundance of the top 5 most abundant TCR lineages prior to ICB (top) and concurrent pool of nonspecific stimulus (bottom). Each line represents a tumor. Abundances are normalized to the pretreatment baseline average of all 100 simulated tumors in the cohort. D. Spearman’s ρ rank correlation between pre-treatment immunogenicity of the rank 1 most immunogenic neoantigen prior to treatment and the extent of immunoediting after one year of ICB treatment. Immunoediting was determined by calculating the fold change of the number of cells expressing the rank 1 neoantigen between the first and last day of ICB. E. Spearman’s ρ rank correlation between pre-treatment clonality of the rank 1 most immunogenic neoantigen prior to treatment and the extent of immunoediting after one year of ICB treatment. F. Spearman’s ρ rank correlation between pre-treatment clonality-weighted immunogenicity of the rank 1 most immunogenic neoantigen prior to treatment and the extent of immunoediting after one year of ICB treatment.

Finally, we evaluated the potential for immunoediting against the rank 1 neoantigen during treatment. Immunoediting is the phenomenon by which cell lineages bearing certain neoantigens are selected against and killed by CTLs, resulting in diminishment or even eradication of certain neoantigens from the immunopeptidome [6]. Across the simulated tumors, we observed no correlation between the immunogenicity nor clonality-weighted immunogenicity of the rank 1 neoantigen and on-treatment immunoediting. There was, however, a very modest negative correlation between the immunoediting and the clonality of the rank 1 neoantigen alone (Figure 6D-F). This suggests that clonality may supersede immunogenicity in provoking lineage-specific selection from CTLs; however, given the very modest effect and borderline p-value, further work in larger patient populations – simulated or otherwise – is needed.

### Model enables exploration of diverse tumor immunopeptidomes

To evaluate model generalizability, RCC-like and MSI-like immunopeptidomes with adjusted tumor growth and mutation rates were simulated and subjected to ICB (Figure 7A-B) [37], [41]. As expected, lower tumor growth and mutation rates led to lower neoantigen burden in RCC-like tumors (Figure 7B). This resulted in poorer predicted responses in these tumors when compared to NSCLC-like and MSI-like tumors, in alignment with clinical experience (Figure 7C) [36], [42], [43]. Even in these neoantigen-poor tumors, however, clonal neoantigen burden correlated with response depth (Figure 7D).

**Figure 7.**
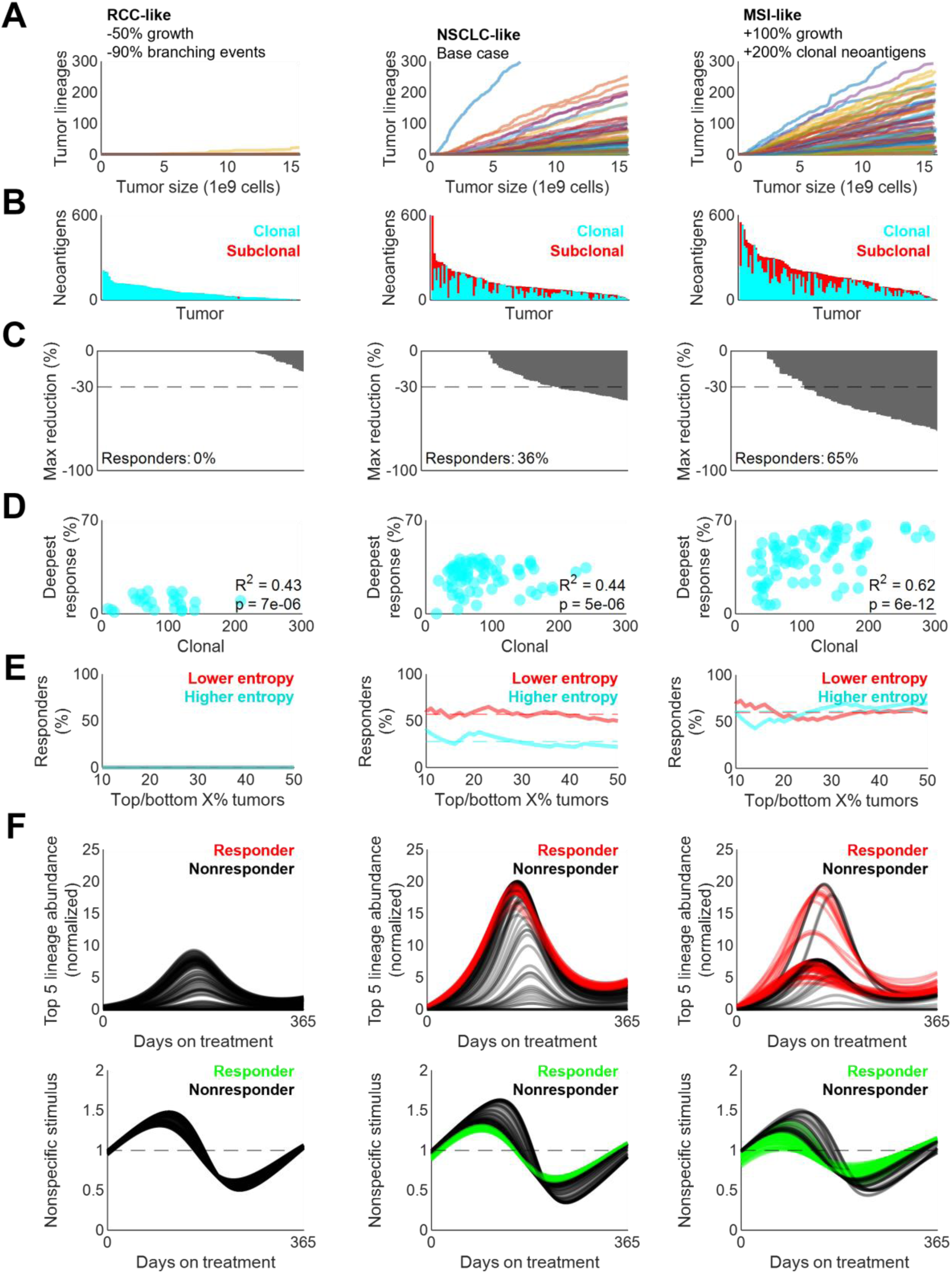
Model describes diverse tumor immunopeptidomes and their response to treatment A. Emergence of unique tumor lineages over the course of simulated tumor growth in RCC-like (left), NSCLC-like (center), and MSI-like (right) tumors. B. Simulated clonal and subclonal neoantigen burden of tumors simulated in (A). Each bar corresponds to a tumor C. Maximum shrinkage of tumors simulated in (A) under treatment constituting 10-fold increases in sensitivity to antigen stimulus and systemic carrying capacity; each bar corresponds to a tumor. The dashed line at corresponds to the 30% or greater tumor shrinkage required to classify as partial response per RECIST 1.1 criteria [35]. D. Pearson’s ρ correlation between response and pretreatment clonal neoantigen burden of tumors simulated in (A). E. Fraction of responses among the X% most and least entropic immunopeptidomes of tumors simulated in (A), with X being indicated by the X axis value. Entropy refers to Shannon entropy. Dashed lines, means. F. Summed abundance of the top 5 most abundant TCR lineages prior to ICB (top row) and concurrent pool of nonspecific stimulus (bottom row) of the tumors simulated in (A). Each line represents a tumor. Abundances are normalized to the pretreatment baseline average of all 100 simulated tumors in the cohort.

This was also the case for neoantigen-rich MSI-like tumors. However, despite their higher neoantigen burden, the most heterogeneous MSI-like tumors were less likely to respond than most homogenous tumors (Figure 7E). Furthermore, the CTL lineages that were most abundant prior to treatment in MSI-like tumors expanded less profoundly upon ICB treatment (Figure 7F). This is in sharp contrast to NSCLC-like tumors, in which oligoclonal CTL expansion was still observed even in nonresponders. The nonlinear relationship between CTL expansion and neoantigen burden warrants further investigation.

## DISCUSSION

Characterizations of the quantitative relationship between the immunopeptidome and CTL clonal dynamics remain lacking. This work provides a mathematical, predator-prey-like framework capable of modeling the co-evolution of the tumor immunopeptidome and TCR repertoire. Built initially upon mouse data, this framework was straightforward to adapt for human tumor simulations. This allowed us to interrogate several clinically observed phenomena, including both pre- and on-treatment changes to the immunopeptidome and TCR repertoire.

While several noteworthy computational models of neoantigen evolution have been published these models consider either a fixed cancer cell population size or calculate death rates as a function of the cumulative immunogenicity of neoantigens expressed by a lineage or clone [2], [15], [44], [45]. No work, to our knowledge, has modeled a growing tumor with dynamic immunopeptidomes and CTL populations. Our results support the directive that neoantigen discovery methods should focus not only on immunogenicity, but also on clonality [46].

One key insight from this work is the shift away from neoantigen “dilution” as the predominant sequelae of intratumoral heterogeneity and toward CTL “distraction”. Several studies report the ability of subclonally expressed neoantigens to dilute an otherwise strongly immunogenic neoantigen; that is, to shield it from immunoediting and dampen anti-tumor killing [8], [14]. Our model supports the importance of neoantigen dilution, showing that neoantigen clonality may be more important than immunogenicity in predicting immunoediting and conceiving biomarkers for responders to ICB. Considering clonality and immunogenicity in tandem is especially informative to ICB response (Figure 6E). However, while these studies and mechanisms are seminally informative to this work, they largely ignore the CTL compartment. Holistically characterizing the anti-immunopeptidome response requires concomitantly assessing how the TCR repertoire evolves across space, time, and tumor types.

Here, we propose the competition for nonspecific stimulus by different CTL lineages as a mechanism for ICB resistance and provide a mathematical modeling framework for its investigation. Our simulations support interclonal CTL competition as a potential reason why highly heterogeneous, subclonal tumors are associated with poor outcomes. We show that CTL diversification during treatment correlates with shallower responses, and that responders who do not diversify have more replete pools of nonspecific stimulus. In addition, many responding MSI-like tumors that responded to ICB have noticeably lower tumor-specific CTL expansion despite greater clonal neoantigen burden. Given these findings, “neoantigen dilution” may be a slight mischaracterization that diverts attention from the fact that CTLs responding to neoantigens are the primary mediators of tumor debulking.

Limitations of this work include its spatially ignorant modeling of metastatic disease as one lumped tumor. Neoantigens can differ between patients and among metastases, potentially leading to increased systemic heterogeneity but decreased local heterogeneity [47]. Moreover, whether the immunopeptidome of individual metastases influences the already spatially biased trafficking of CTLs remains an open question. Are neoantigen-reactive CTLs more likely to linger in tumors where their target neoantigen is present? What if the neoantigen is highly subclonal – what is the probability they will physically encounter it? Is there a critical clonality threshold? These are critical questions that the model is not currently equipped to address.

Our work is primarily informed by data from two mouse models and two NSCLC studies. While the role of neoantigen clonality in NSCLC is generally consistent with its role in melanoma and colorectal cancer, it could differ in other tumor types such as low-grade glioblastoma and prostate cancer [37]. Moreover, the simulated response rate in MSI-like tumors was above that observed in the clinic, suggesting variability in treatment or CTL potency across tumor types.

Ongoing clinical studies profiling the effect of ICB on TCR repertoire diversity in CRC may shed additional light on the veracity of our observations [48]. More generally, studies considering the coevolution of the cancer immunopeptidome and TCR repertoire diversity (or lack thereof) will be critical to achieving long-term disease control and cure.

In summary, this predator-prey-like framework accurately described clinically observed immunopeptidome diversity and recapitulated clinical evidence supporting the correlation between clonal neoantigen burden and ICB response. By including a dynamic CTL compartment, the model could also describe the effects of TCR diversity on predicted responses. Analysis of the tumor-immune axis also analyses also suggested that neoantigen immunoediting may correlate more strongly with pretreatment clonality than immunogenicity. This model can be readily adapted to future data published in other tumor types through simple, biologically motivated adjustments to parameter values. Further refinement could facilitate low-cost theoretical explorations of histology-specific therapeutic strategies in advance of developing and deploying them in the clinic. This work supports the further development of predator-prey-like models to elucidate nonlinear tumor-immune dynamics for clinical translation and utility.

## DECLARATIONS

### Patient consent for publication

Not applicable.

### Ethics approval

Not applicable.

### Availability of data and material

All original code has been deposited at Zenodo (10.5281/zenodo.6596442) and is publicly available. All data collected for simulation benchmarking have been made available in the Supplement.

### Authors’ contributions

Conceptualization, T.Q. and Y.C.; Methodology, T.Q.; Software, T.Q.; Validation, T.Q.; Formal Analysis, T.Q.; Investigation, T.Q.; Resources, Y.C.; Data Curation, T.Q.; Writing – Original Draft, T.Q.; Writing – Review & Editing – T.Q., Y.C., and B.V.; Visualization – T.Q.; Supervision – Y.C.; Project Administration – T.Q. and Y.C.; Funding Acquisition – Y.C.

## Acknowledgements

The authors thank Amber Cipriani and J. Justin Milner for their helpful input during study design and manuscript preparation.

## SUPPLEMENTAL MATERIAL

## SUPPLEMENTAL TABLES AND FIGURES

**Table S1.** Model parameters

**Figure S1.**
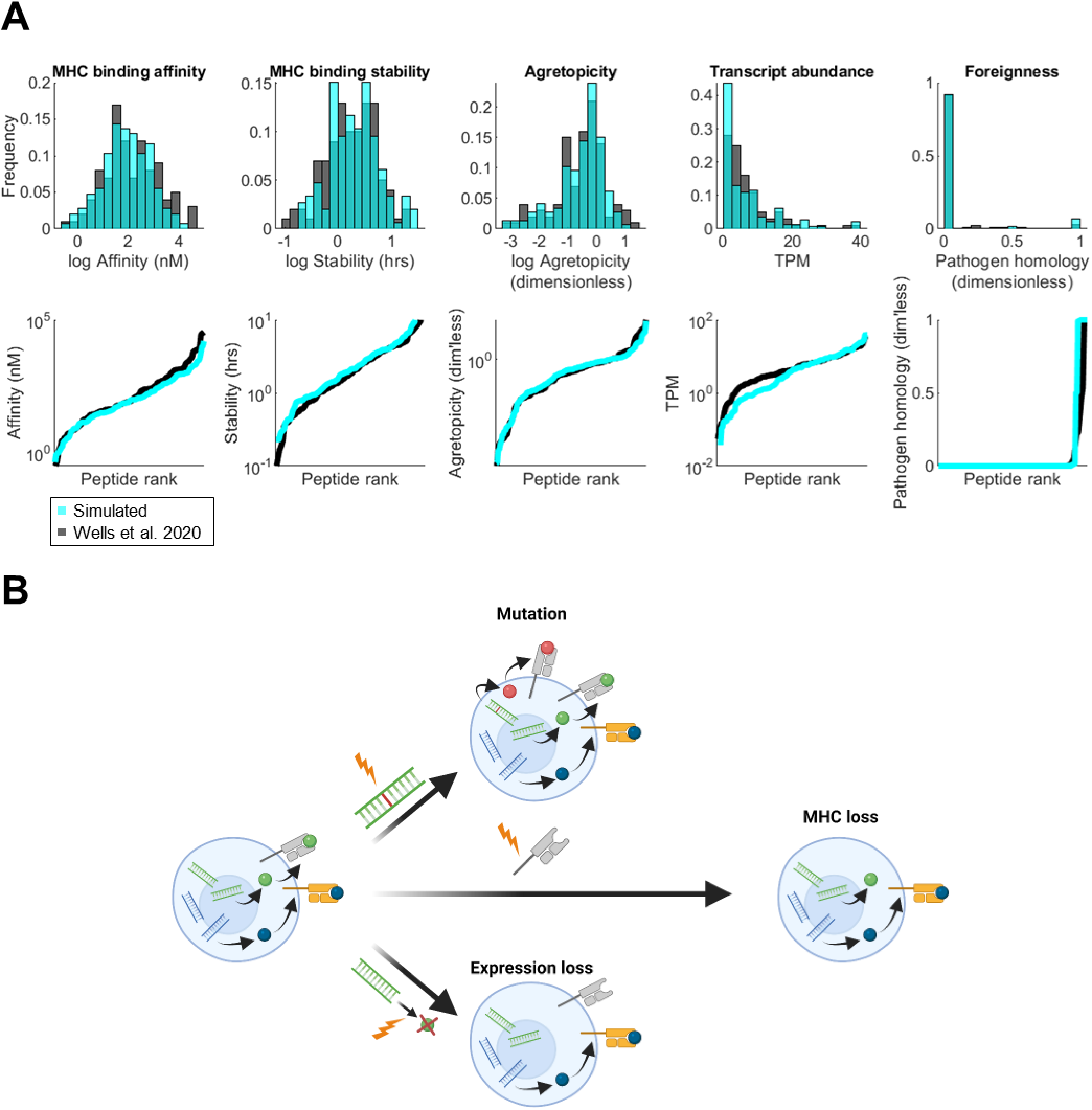
Model-simulated neoantigens account for stochastic events and TESLA criteria A. Each neoantigen was assigned values for the TESLA parameters drawn from the distributions for NSCLC tumors by Wells et al. in [4]. TPM, transcripts per million. Distributions (top) and values (bottom) of sampled and experimental parameters from 146 NSCLC-derived peptides are shown. B. Schematic of stochastic neoantigen events. A founder cell is established with a random number of neoantigens and assigned a neoantigen gain rate from a lognormal distribution. Each neoantigen was also assigned to be presented by one of three MHC alleles. Neoantigen and MHC expression could also be lost by a cell during tumor growth.

**Figure S2.**
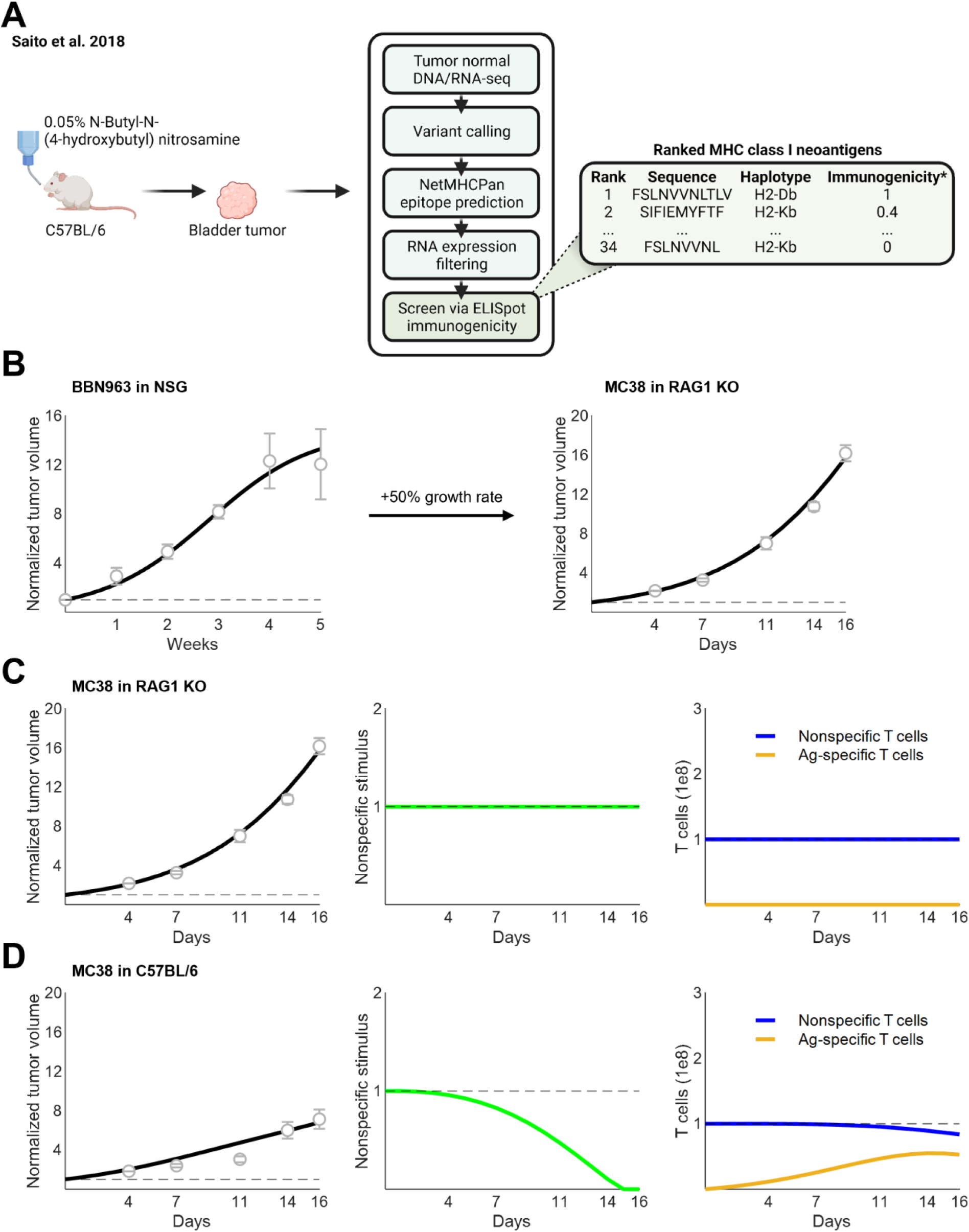
Characterization of the BBN963 immunopeptidome and validation in MC38 cells A. BBN963 cells were established by chronic exposure of C57BL/6 mice to 0.05% N-Butyl-N-(4-hydroxybutyl) nitrosamine. Neoantigens were called and screened for bona fide immunogenicity in an ex vivo ELISpot assay. Additional details can be found in [23]. B. Growth dynamics of MC38 were simulated using the same parameters as BBN963 cells, except growth rate, which was increased 50% [25]. C. Simulated tumor volumes (left, black) normalized to baseline volume (dashed gray line) compared against growth dynamics in RAG1 KO mice reported by Efremova et al. (gray circles and error bars). Nonspecific stimulus (center) and CTL populations (right) are shown. Dashed lines, 1. D. Simulated tumor volumes (left, black) normalized to baseline volume (dashed gray line) compared against growth dynamics in C57BL/6 mice reported by Efremova et al. (gray circles and error bars). Nonspecific stimulus (center) and CTL populations (right) are shown. Dashed lines, 1.

**Figure S3.**
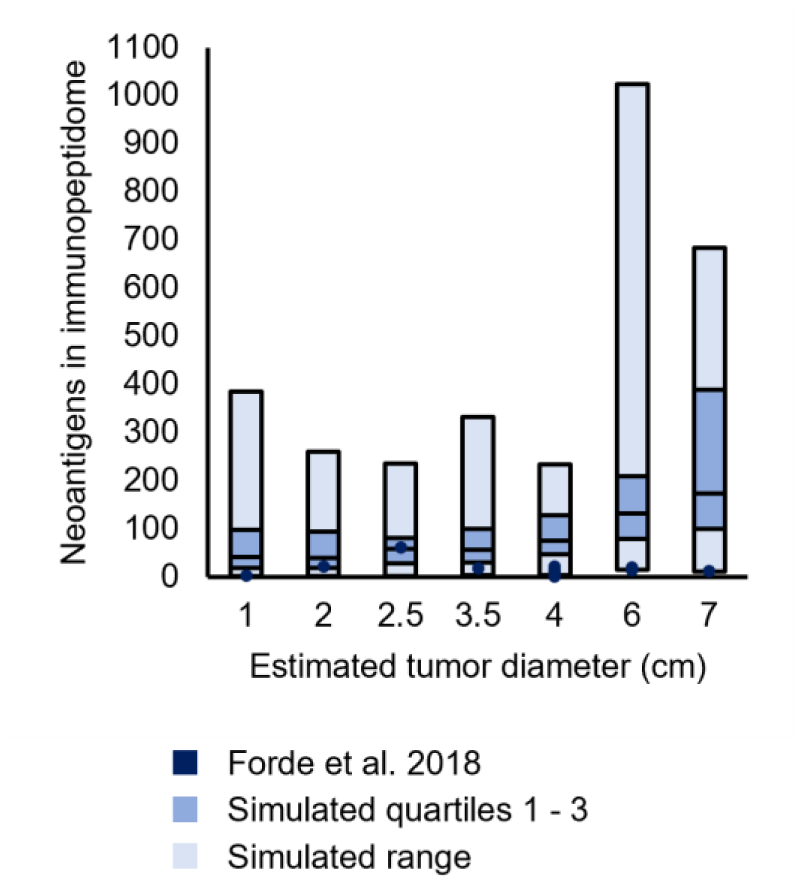
Simulated neoantigen burdens were within range of an independent cohort A. Neoantigen predictions reported by Forde et al. 2018 in 12 patients with sufficient pretreatment tissue to characterize candidate neoantigens were assessed. Total neoantigens were calculated by filtering positive netCTL classification. To apply Wells’ criteria, we calculated agretopicity by taking the ratio of somatic and wild-type MHC binding affinity. For predictions containing somatic peptides of identical sequences but different HLA alleles, the stronger predicted binding was used. Binding stabilities were calculated with NetMHCStabPan using default parameters. As biopsies were single-region, neoantigen clonality was not assessed. Tumor diameters were estimated for each tumor using conventional NSCLC staging criteria. For each of seven discrete estimated tumor diameters, a cohort of 40 tumors was simulated.

**Figure S4.**
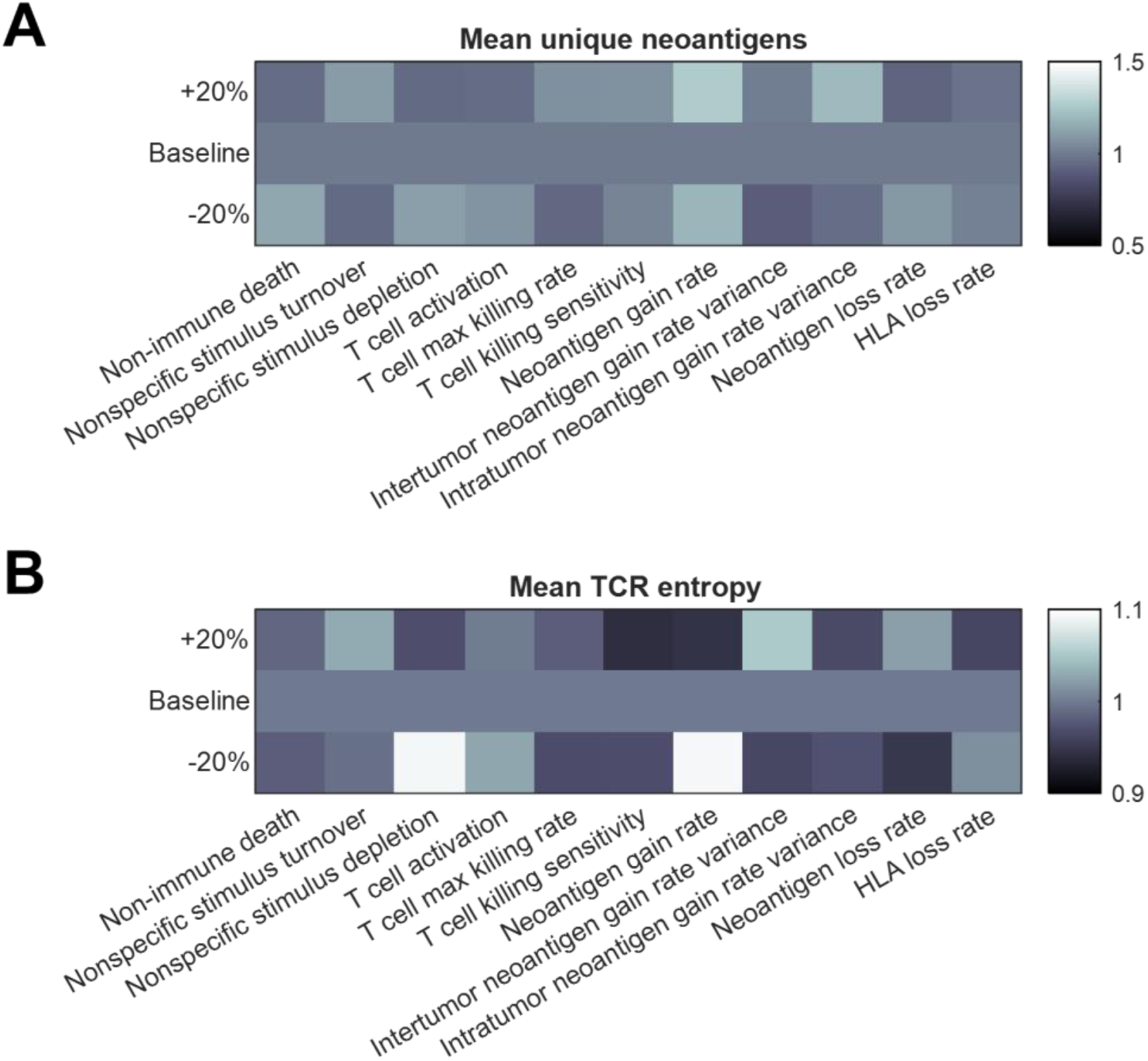
Local sensitivity analysis of parameters indicates model robustness A. Mean number of unique relative to baseline (N=100 simulations per condition). No significant fold differences in neoantigen number were found from pairwise Student’s t-tests with neither nominal nor Bonferroni-corrected significance thresholds. B. Mean TCR entropy relative to baseline (N=100 simulations per condition). No significant fold differences in Shannon entropy were found from pairwise Student’s t-tests with neither nominal nor Bonferroni-corrected significance thresholds.

